## Supplementary figures and images for "Cardiomyocyte apoptosis contributes to contractile dysfunction in stem cell model of *MYH7* E848G hypertrophic cardiomyopathy"

### Supplemental Figure 1

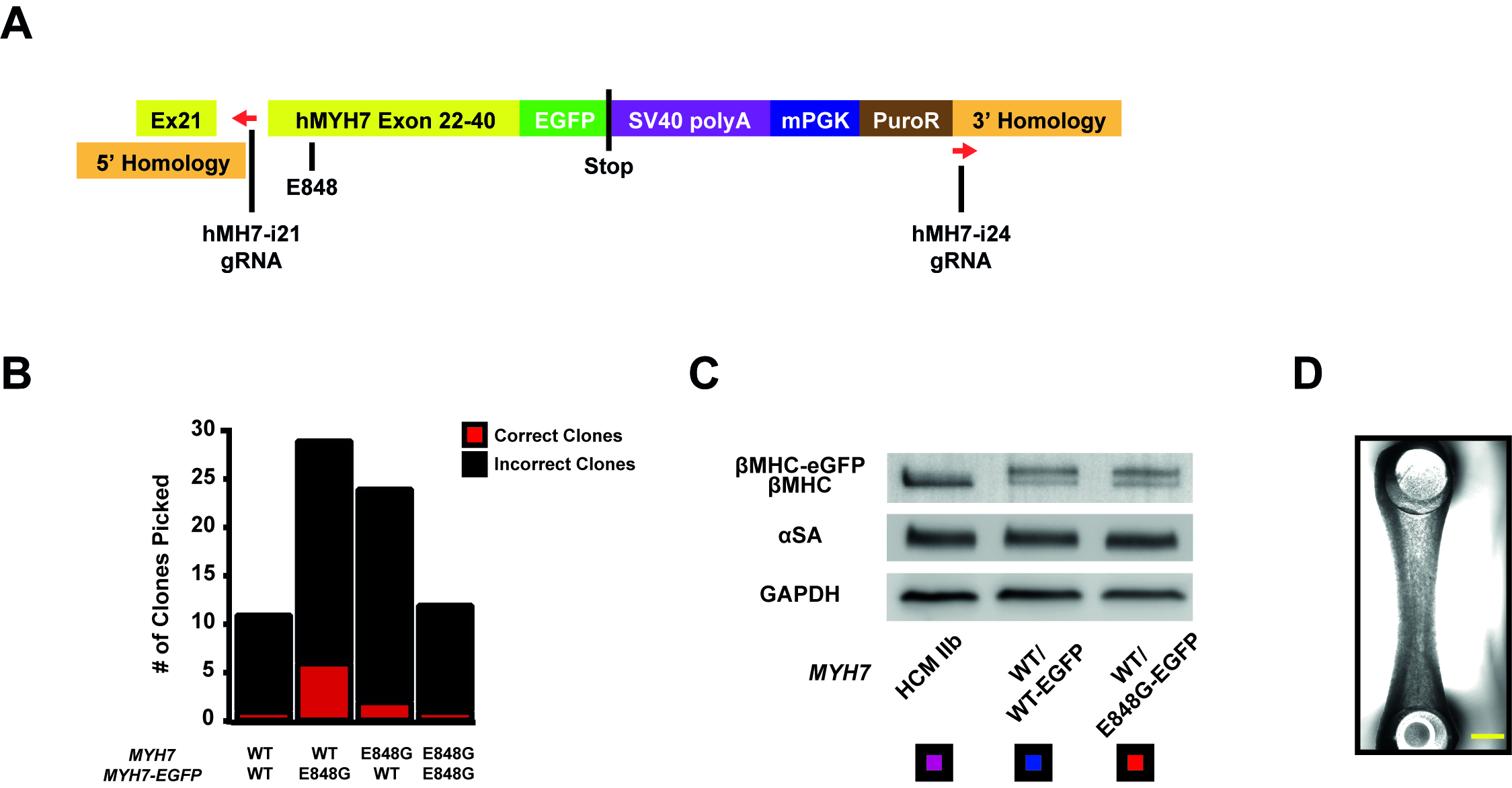

### Supplemental Figure 2

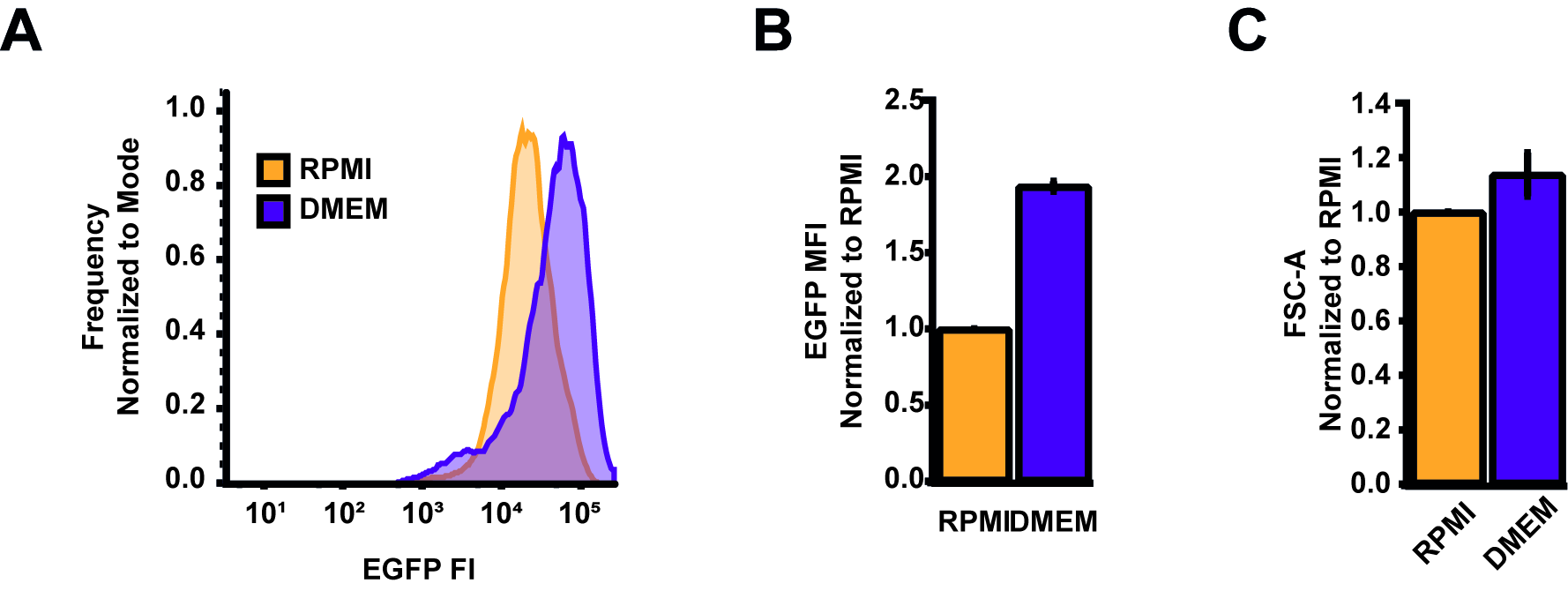

### Supplemental Figure 3

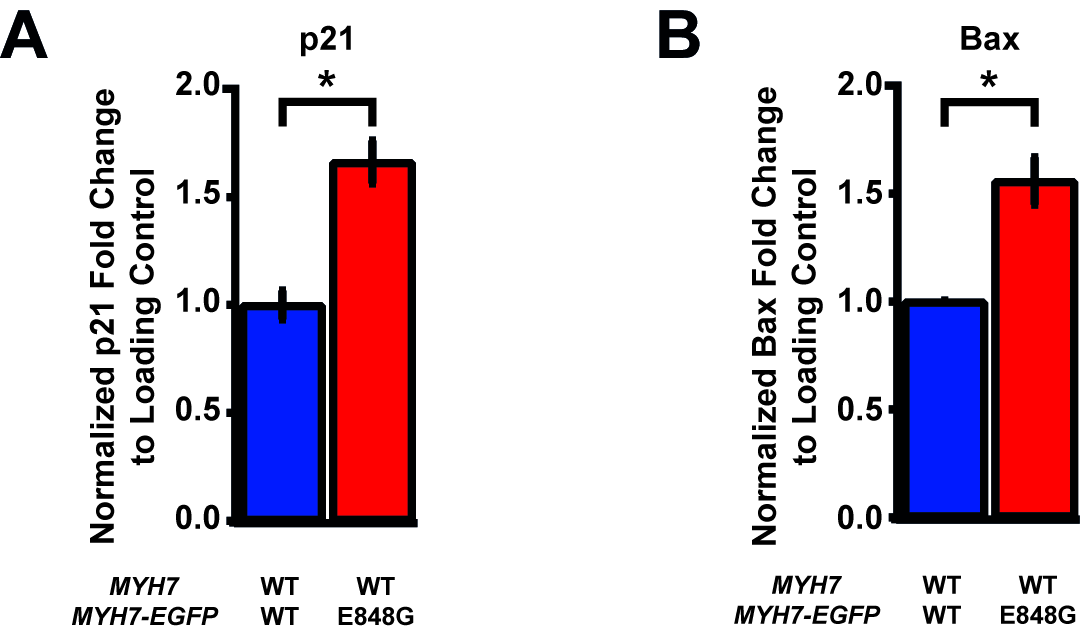

### Supplemental Figure 4

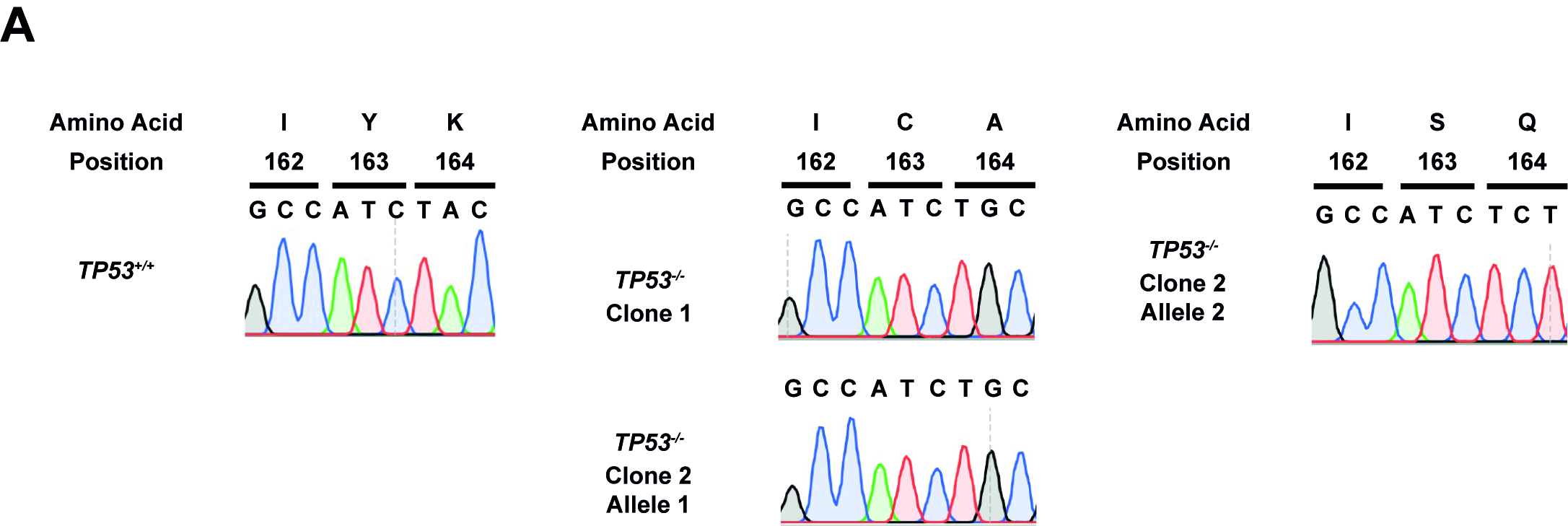
